# Goals as dynamical attractors: a momentum-based account of stable and flexible goal commitment

**DOI:** 10.64898/2026.05.06.723407

**Authors:** Sneha Aenugu

## Abstract

Human goal pursuit is often marked by persistent activity toward achieving an objective, as well as flexibility in switching objectives based on environmental demands. How humans balance the stability and flexibility necessary for goal pursuit is the key question of this study. We propose that goal pursuit generates dynamic attractor modes in policy landscapes that produce stability in goal pursuit. The attractor properties are modulated through progress monitoring, allowing for the flexibility necessary to switch objectives in favor of alternative goals. Through simulations and behavioral cloning of human participants performing an extended goal selection task, we show how dynamic modes can develop in the latent spaces of recurrent neural networks trained with reinforcement learning. We develop metrics to quantitatively assess the attractor qualities of dynamic modes, validating them against synthetically generated dynamical systems, and use them to investigate the context modulation of attractor modes during goal pursuit. We then proceed to develop a circuit-level account of goal persistence incorporating self-excitation and cross-inhibition as motifs for fast, self-sustaining dynamics modulated by slow, progress-integrating momentum and context signals. Lastly, we show that the switching costs experienced while managing multiple goals are an emergent property of resistance to the intrinsic dynamics of goal pursuit, thereby contributing a fresh perspective on the dynamics of extended goal pursuit.

## 1 Introduction

We pursue multiple goals concurrently in our daily lives. Most often, however, we pursue one goal at any specific instance in time and maintain persistent activity in its favor. Say, writing a section of a manuscript requires you to sit at your desk and persistently engage with it. However, an email ping distracts you, and you switch over to the new task of checking your messages. We constantly manage multiple such goals in our daily lives, which requires us to maintain stable goal states for progressing in a task while also remaining flexible enough to switch over to alternative rewarding tasks in the environment. This fundamental aspect of intelligence is essential to all agents aspiring toward general intelligence: the ability to manage diverse tasks under various environmental tradeoffs.

The mechanistic basis of how we maintain stability and flexibility in the pursuit of extended goals is an open question. A few theories of goal persistence address the stability aspect of the question by invoking resource-rational accounts of decision-making. Goal persistence is thought to reduce policy complexity, enabling us to perform tasks by avoiding intensive computations. Some models of goal persistence incorporate policy bias or perseveration terms as inductive biases for self-sustaining activity in favor of goal pursuit. On the other hand, flexibility is thought to be a product of reinforcement learning of reward rates in the environment. However, as most extended goals deliver rewards only after goal completion, goal progress can be thought of as a pseudo-reward indicating the prospects of resolving a given goal [1]. [2, 3] proposed a momentum model of goal selection, where goal progress and its derivative (the rate of goal progress) can jointly influence decision-making, and their product, goal momentum, can serve as the drive influencing goal selection. This model was shown to capture several behavioral properties in humans pursuing extended goals: human overpersistence patterns, preference for goal progress, and increased rates of revisiting goals with greater progress after switching away from them. Such models remain good descriptive accounts of goal selection; however, they fall short of providing a mechanistic explanation of their basis in the brain, specifically how they enable the maintenance of stable goal states robust to environmental distractors and dynamic states that can be perturbed through rewards in the environment [4, 5].

The theory of dynamical systems holds great promise for generating a mechanistic explanation of extended goal pursuit, especially regarding how stability-flexibility tradeoffs are realized [6–8]. Attractor properties developed by dynamical networks are a strong analogue for architectures that can generate and maintain stable, self-sustaining intrinsic dynamics while still retaining flexibility by being pushed around by input-driven dynamics [9]. [10] made that connection by framing goals as far-from-equilibrium attractors of a dynamical system. [11] proposed an attractor-dynamics account of resolving motivational conflict by viewing behavioral switching either as a bifurcation or as a stochastic escape from an attractor basin. Attractor models of goal pursuit thus translate the metaphorical insight of persistence as a pull toward specific goals into a rigorous quantitative interpretation of decision-making as traversal through an energy landscape guided by context and rewards in the environment.

Recurrent neural networks (RNNs) are a strong test bed for investigating dynamic modes of human decision-making, given their ability to integrate past experiences into a latent context that translates into action policies. Studying the dynamics of the latent space can therefore lend us an understanding of the behavioral dynamics seen in human decision patterns [12]. [13] put forth a modeling approach that employs tiny RNNs to discover cognitive strategies by fitting RNNs to human choices and showed that it can be a viable predictive model of human behavior. In this study, we use RNNs as both normative and descriptive accounts of human decision dynamics. We first simulate RNNs to investigate conditions under which they emulate human persistence patterns, and later we clone human behavior with RNNs to describe the dynamic modes underlying human persistence patterns.

While RNNs can be descriptive of human dynamic modes and serve as a baseline model of decision dynamics, they tell us little about the mechanistic implementation of such modes in the brain. Specific network motifs combining excitatory and inhibitory firing are known to facilitate self-sustaining behavior [14] and can facilitate selective attention in goal pursuit [15]. [16] showed in a discrete recurrent circuit that a combination of slow excitation and fast inhibition yielded persistent attractor dynamics that enabled stability and rapid switching during fly navigation. In this paper, we use a self-excitation and cross-inhibition network motif to generate fast self-sustaining dynamics that facilitate goal commitment and are dynamically modulated by slow, progress-integrating momentum signals. We investigate how well this model explains human behavior and study its dynamic modes as modulated by behavioral context. Through this work, we generate explicit mechanistic hypotheses about how goal states are stably maintained and dynamically shifted that can be tested for circuit-level neural implementation using electrophysiological recordings in animals or humans performing extended goal pursuit tasks.

## 2 Experiment and behavior

We use the extended goal pursuit paradigm introduced in [2] to model human goal selection. In the game, participants are free to pursue any of the three extended goals—collecting car, hat, or cat suits—and are free to switch between them without penalty. To finish a specific suit (goal), participants need to collect seven tokens of the same kind (Figure 1-left). The game proceeds in rounds, and participants can choose to acquire any of the three tokens at a given instance. The success of the actions is probabilistic, and the probabilities vary throughout the task in blocks. Say, in a specific block in the task, cat tokens might be more available—for example, if you choose to acquire the cat token, you will be given the token with an 80% chance and no token with a 20% chance. On the other hand, in the same block, if you choose to acquire the hat token, you can get the token with a 20% chance and fail with an 80% chance. The task has three block types—80-20, 60-40, and 70-30—wherein the 80-20 block has one goal that gives tokens with an 80% chance, while the other two goals give tokens with a 20% chance. Participants need to infer the identity of the goal that has the highest probability within a block for optimal goal pursuit and switch adaptively when the block changes and a new goal becomes the dominant one. The objective of the game is to complete as many goals as possible. We acquired the openly available human behavior dataset from [2] to model the behavior of 44 participants who played the game.

**Figure 1:**
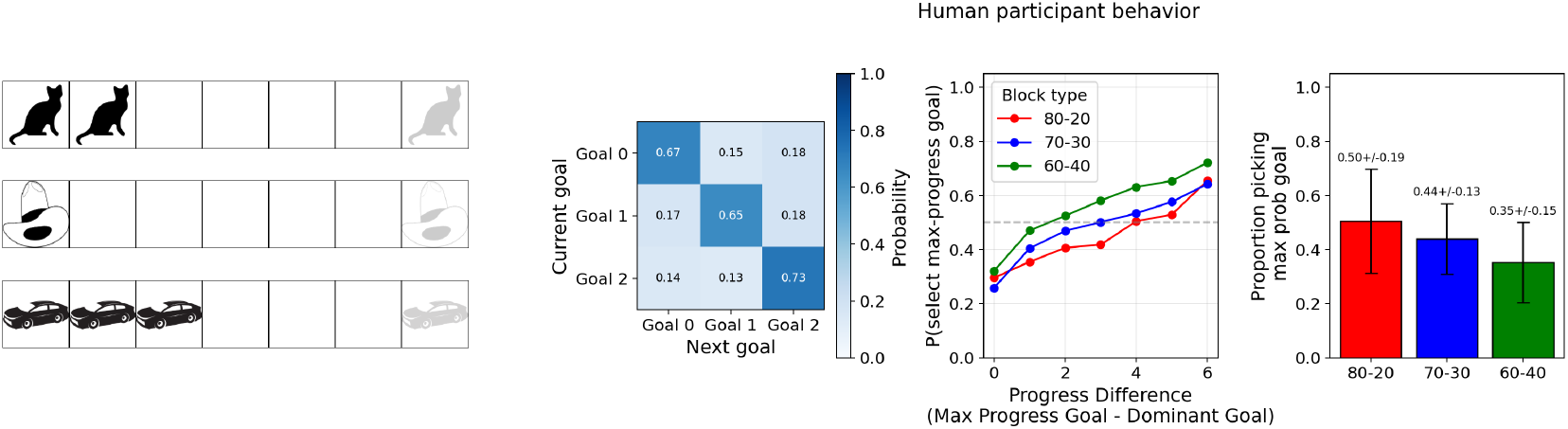
Task and behavior. (left) Extended goal pursuit task: Collect 7 tokens of the same kind to finish a suit and collect rewards. Participants can switch between suits (cat/ hat/ car) without penalty. (right) Human behavior in the task: (Panel 1) Probability of transitioning from one goal to the next across adjacent rounds. (Panel 2) Participants goal selection probability plotted against the progress difference between the goal with the maximum progress — max-progress goal — and the goal which gives tokens with a higher chance — max-prob / dominant goal. (Panel 3) Participants probability of selecting the max-prob goal in different block types.

Participants in the game maintain persistent goal states for extended durations. Participants’ aggregate transition probability matrix (Figure 1) shows a significantly higher probability of staying with the same goal across adjacent rounds. Participants also show a preference for progress, wherein their probability of selecting a goal increases with progress toward that goal. Figure 1 shows participants’ goal selection probability plotted against the progress difference between the goal with the maximum progress (max-progress goal) and the goal that gives tokens with a higher probability (max-prob goal). Participants also show sensitivity to block types. In the 80-20 block, where there is a larger incentive to pick the max-prob goal, participants show a lower preference for progress compared to the other blocks. Humans in the game were shown to overpersist beyond what is optimal for the task. [2] constructed algorithms that beat human performance (measured by the number of goals completed) by a significant margin. One of the agents prospectively estimates the time to goal completion by learning the probability structure of goals within the block while discounting future rewards. This agent shows progress preference and persistence in the task, yet substantially outperforms humans, who show patterns of overpersistence compared to the prospective agent. We now train recurrent neural networks to play the same game as participants to investigate what parameterizations can produce persistence trends similar to those observed in humans.

## 3 Modeling the dynamics of extended goal commitment using RNNs

### 3.1 RNNs optimizing for discounted future returns in the task exhibit persistence and preference for progress

We trained a recurrent actor-critic network to play the extended goal pursuit task, optimizing for total rewards in the game. The network takes the following observation vector from the environment:

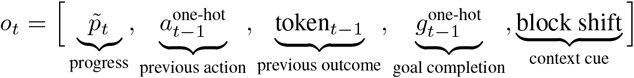

It first embeds the observation and passes it through a gated recurrent unit (GRU), which ensures the latent state integrates past observations and actions.

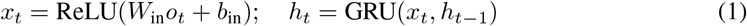

We then define the policy (actor) and value (critic) based of the latent state.

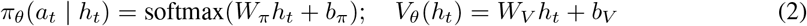

We trained the network to optimize the number of goals finished in the task (details provided in Appendix Section B). The recurrent policy architecture learns to perform the task, albeit without displaying the degree of persistence seen in human participants (see Appendix Figure 4). The network also does not show the preference toward progress observed in humans. We could modulate the network’s persistence levels by tweaking the variable that discounts future returns in the game. It is widely documented that humans incorporate discounting in their valuation [17], which partly accounts for their persistence with the max-progress goal. We therefore modified the network’s objective to optimize discounted rewards in the game (*γ* = 0.9), which increased persistence levels to those of human participants and also reproduced the progress preference seen in humans (Appendix Figure 4). We then let the trained network play the game and plotted the principal components (PCs) of the latent-space trajectory over time. Furthermore, the latent space shows a clear gradation of goal progress and goal progress rate, indicating that the network’s latent space adequately represents key factors necessary for playing the game. We next clone participants’ behavior using recurrent neural networks and analyze their latent space for dynamic modes of human persistence.

### 3.2 Behavior cloning through recurrent neural networks reveals attractor-like modes in goal-directed policy dynamics

We model each subject’s sequential decisions using a recurrent neural network (RNN) trained via behavioral cloning [18, 19]. At each trial *t*, the model receives an observation vector *o*_*t*_ encoding current goal progress, previous action, previous reward outcome, previous goal completion, and block-boundary signals.

The RNN updates a latent state *h*_*t*_ and produces action probabilities:

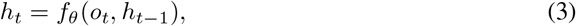

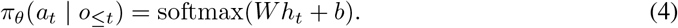

We fit the parameters independently for each subject by minimizing the negative log-likelihood of observed actions:

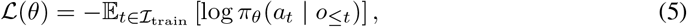

using an interspersed train/validation/test split (75/12.5/12.5). We evaluated performance using held-out negative log-likelihood, prediction accuracy, and agreement with empirical choice frequencies. We provide further details on the behavioral cloning procedure in Appendix Section C. The fitted hidden states {*h*_*t*_} are used for downstream dynamical analyses.

#### Context-Dependent Intrinsic Dynamics

To characterize the internal dynamics of the trained recurrent neural networks, we analyze their intrinsic velocity fields in a low-dimensional latent space. We first extract the hidden-state trajectory 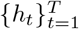 for each subject and project it onto the top principal components using PCA. Let *z*_*t*_ ∈ ℝ^2^ denote the projection onto a chosen 2D plane. To define context-conditioned dynamics, we group time points according to the previous goal and current progress level. For each context *c*, we construct a representative observation *ō*^(*c*)^ by averaging observations within that context. Rather than evaluating dynamics only along observed trajectories, we compute the intrinsic flow field over an artificial grid of latent states spanning the PCA space. Each grid point is mapped back to a corresponding hidden state, with non-visualized principal components fixed to their mean values. Holding the context fixed via *ō*^(*c*)^, we evaluate a single-step RNN transition: *h*^′^ = *f*_*θ*_(*ō*^(*c*)^, *h*) and define the intrinsic velocity as *v*(*h*; *c*) = *h*^′^ − *h*. This velocity is then projected into the PCA plane to obtain a vector field over the grid.

To identify candidate attractors, we locate regions of low velocity magnitude in the projected field and extract local minima of |*v*(*h*; *c*) | , which we interpret as slow points—proxies for fixed points in the underlying high-dimensional dynamics [20]. The resulting vector fields, evaluated on a grid in PCA space and visualized as quiver plots, are shown in Appendix Figure 5. This procedure (elaborated further in Appendix Section C.1) yields context-dependent dynamical landscapes that reveal how goal representations are stabilized or destabilized as a function of behavioral state. Qualitatively, the flow fields exhibit point-attractor-like structure, with trajectories converging toward compact regions in latent space. While they might not qualify as strict fixed-point attractors, these quasi-attractor-state properties are modulated by increasing goal commitment [21]. To quantify this effect, we next introduce metrics that capture the “pointiness” of these quasi-attractor modes.

#### Quantifying Point-Attractor Structure

To quantify the extent to which learned dynamics exhibit point-attractor structure, we analyze intrinsic velocity fields within each behavioral context (previous goal × progress level). Candidate fixed points are identified as regions of low velocity magnitude.

For each candidate, we evaluate whether it satisfies the defining properties of a true point attractor using a strict composite metric. Specifically, we define a *strict point-attractor score* that combines multiple necessary dynamical criteria. This score is high only when a candidate exhibits inward flow, basin support, and full local stability. The components are:

- **Inward alignment:** Measures whether nearby vectors in the field point toward the candidate. High values indicate that the local flow is directed inward, consistent with attractor dynamics.
- **Point-specific stability:** Assesses local stability using the Jacobian of the vector field. The score penalizes both unstable directions (positive eigenvalues) and neutral directions (near-zero eigenvalues), ensuring that only fully stable fixed points receive high values.
- **Basin support:** Quantifies whether nearby trajectories converge toward the candidate over short rollouts. This captures whether the candidate has a meaningful basin of attraction beyond its immediate neighborhood.

These components are combined multiplicatively to yield a strict point-attractor score, such that a high score requires all properties to be simultaneously satisfied. We provide details of the metric and its validation using synthetic dynamical systems in Appendix Section D. We compute these metrics separately for flow fields separated into low- and high-progress contexts and compare them within subjects using paired statistical tests. Figure 2 (bottom) shows that high-progress states exhibit significantly higher strict point-attractor scores than low-progress states, consistent with the idea that increasing progress stabilizes goal representations by deepening attractor dynamics in latent space. While behavioral cloning of human goal selections provides insight into the context-dependent intrinsic dynamics of choice patterns, we still lack a mechanistic account of goal persistence that can point to specific circuit motifs resulting in the dynamic modes apparent in human behavior. In the next section, we formulate a circuit-level interpretable dynamical-systems account of human goal pursuit.

**Figure 2:**
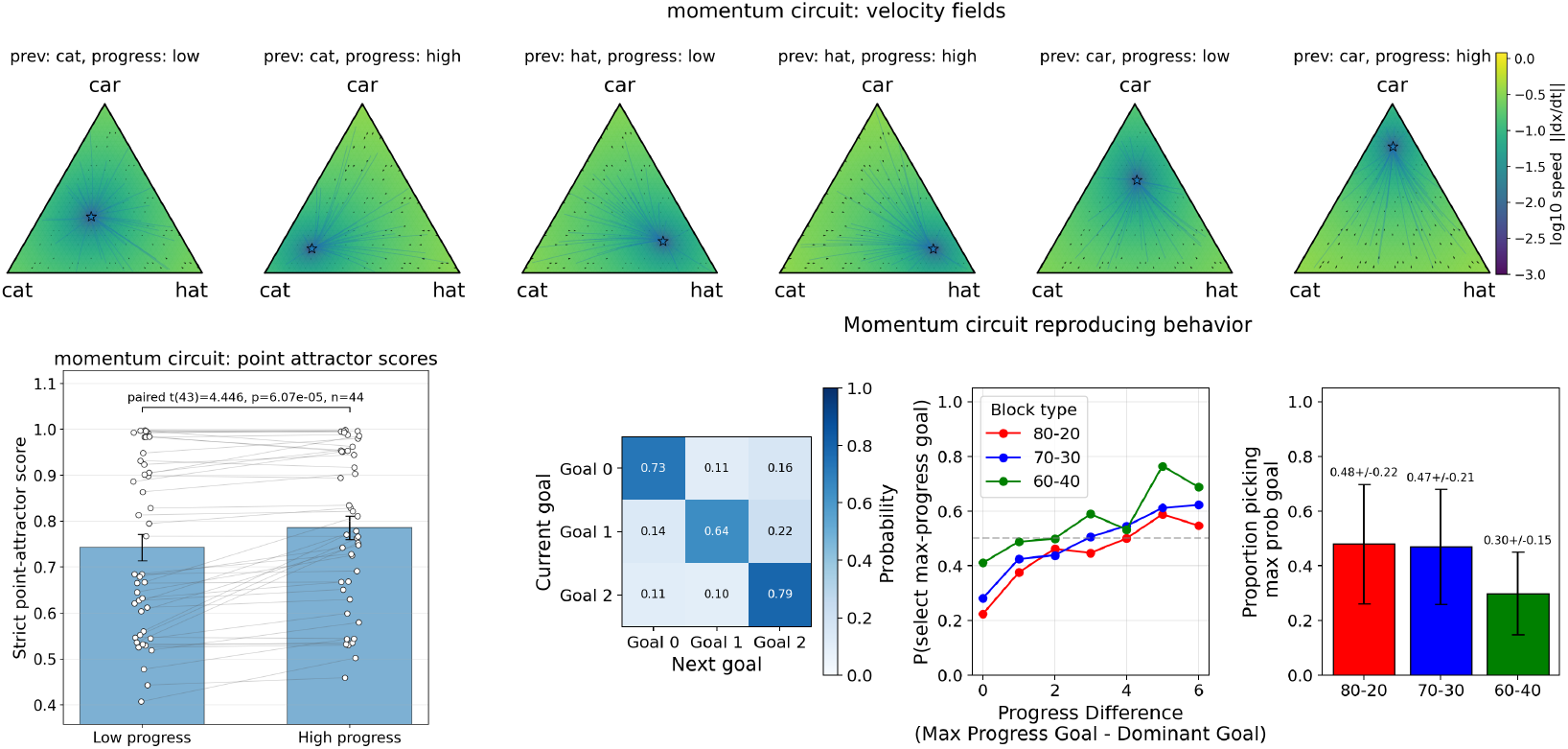
(top) Context-dependent intrinsic velocity fields of the momentum-circuit model of a single participant in the simplex space showing point-attractor-like structure. Each panel shows the velocity field for a specific previous goal and progress level. (bottom) (left) Significantly higher point-attactor scores in high-progress contexts compared to low-progress contexts showing attractor-structure evolution with environmental context. (right) Momentum-circuit model reproducing human persistence patterns and progress preferences.

## 4 A momentum-based circuit model of goal pursuit

We introduce a momentum-based circuit-level implementation of goal selection, in which goal commitment emerges from recurrent competition in policy space. Persistence arises from fast competitive dynamics between goal states, modulated by progress-dependent signals, environmental context, recent choice history, and feedback from the selected action.

We represent goal commitment as a distribution over three goals, **x**_*t*_ ∈ Δ^2^, where the 2-simplex is defined as:

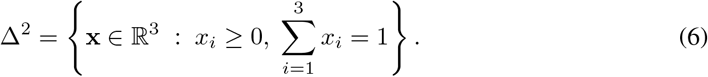

### Drive and policy

At each time step, the circuit computes a drive vector:

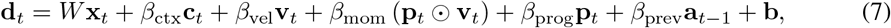

where *W* ∈ ℝ^3×3^ encodes recurrent competition, **c**_*t*_ is a context estimate, **p**_*t*_ ∈ [0, 1]^3^ is goal progress before choice, **v**_*t*_ is an asymmetric temporally integrated progress velocity, **a**_*t* −1_ is a one-hot vector encoding the previous action, **b** is a goal-specific bias, and ⊙ denotes elementwise multiplication. Progress therefore enters both directly through *β*_prog_**p**_*t*_ and indirectly through velocity and progress-gated momentum.

### Recurrent competition

The recurrent interaction matrix implements self-excitation and cross-inhibition between goal states [22].

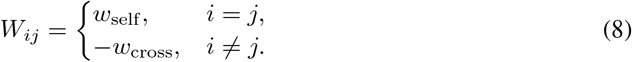

Equivalently, the recurrent contribution can be written componentwise as

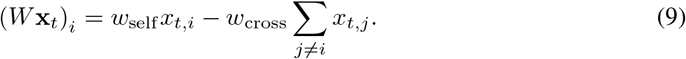

### Asymmetric momentum computation

Humans pursuing goals care not only about how much progress they have made, but also about how fast they continue to make progress. To capture this, we define an asymmetric leaky progress-velocity trace. First, instantaneous progress change is computed as

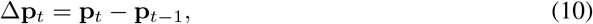

with Δ**p**_0_ = **0**. Positive and negative changes are separated as

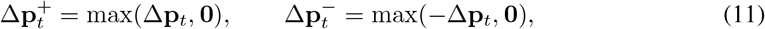

where the maximum is applied elementwise. The velocity trace is then updated according to

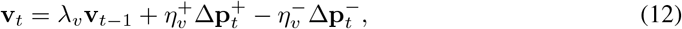

where 0 *< λ*_*v*_ *<* 1 is the velocity decay and 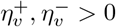 are learned gains for positive and negative progress changes. This allows positive progress and negative/no-progress evidence to exert different effects on the velocity trace. This leaky updating of progress gradients induces a slow-integrating momentum signal [23], akin to momentum in gradient descent updates [24].

Another momentum-like computation that further builds inertia at high-progress levels is the progress-gated velocity signal: **m**_*t*_ = **p**_*t*_ ⊙ **v**_*t*_. Thus, **v**_*t*_ captures a temporally smoothed and valence-asymmetric estimate of recent progress change, while **p**_*t*_ ⊙ **v**_*t*_ amplifies commitment when progress is high and recently increasing.

### Context computation

While slow-integrating momentum signals provide an approximate measure of ongoing goal approach, they do not by themselves encode broader environmental statistics that can inform goal selection, such as block structure in the task. The context vector **c**_*t*_ ∈ ℝ^3^ encodes a running estimate of goal-specific success rates within each block. Let *S*_*t,i*_ and *F*_*t,i*_ denote the number of successes and failures for goal *i* up to time *t*. We compute:

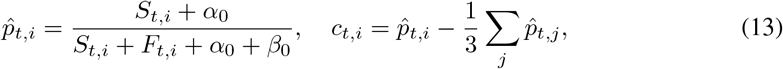

so that **c**_*t*_ reflects the relative advantage of each goal within a given block.

### Action policy

Actions are sampled from a softmax policy, where *τ >* 0 is the softmax temperature.

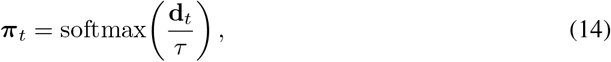

### State dynamics

The latent goal state evolves via leaky integration toward the current policy, with an additional choice-feedback term that pulls the latent state toward the action selected on the current trial. Let **a**_*t*_ denote a one-hot vector encoding the chosen action at time *t*. The unnormalized next state is

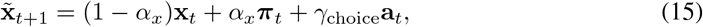

and the simplex-constrained latent state is obtained by rectification and normalization:

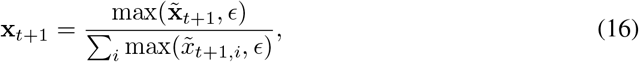

where 0 *< α*_*x*_ *<* 1, *γ*_choice_ ≥ 0, and *ϵ* is a small numerical floor.

The model defines a dynamical system in which recurrent interactions *W* **x**_*t*_ generate attractor structure over goals, while context, direct progress, asymmetric progress velocity, progress-gated momentum, recent action history, and choice feedback modulate this landscape based on ongoing task performance. This interaction gives rise to persistent goal states and switching behavior as emergent properties of the underlying dynamics. We fit the model independently for each subject using maximum likelihood and analyze the resulting dynamics in the simplex policy landscape.

### Simplex Landscape of Goal Dynamics

To visualize the dynamical modes of the fitted circuit model, we construct vector fields over the goal simplex. Given a goal state **x**, the model updates it based on recurrent dynamics and fixed task-dependent inputs, inducing movement within simplex policy space. For a fixed input condition (**c, p, v, a**_prev_) and drive, **d**, defined as per equation (7), define

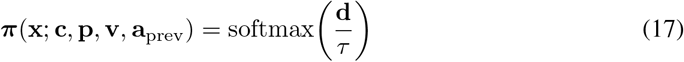

The corresponding policy-driven update direction is

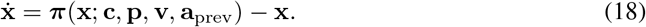

When visualizing the full latent-state update including choice feedback, the vector field can instead be defined using the normalized update map

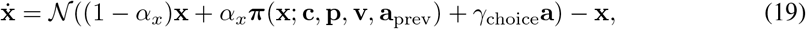

where 𝒩 (**z**) = max(**z**, *ϵ*)*/*Σ_*i*_ max(*z*_*i*_, *ϵ*). In the intrinsic policy-landscape analyses, we fix the behavioral context by holding all non-state inputs constant. Specifically, for each condition, defined by previous goal identity and progress level, we compute representative values of progress, velocity, momentum, previous action, and context signals by averaging over matching trials. These anchor variables define a fixed input condition, allowing us to analyze the induced dynamical system over the goal state alone. We evaluate 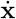 on a dense grid over the simplex and visualize vector directions, speed 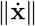 , and simulated trajectories from random initial states. Candidate attractors are identified by simulating trajectories under the vector field and clustering their endpoints. Stable goal states appear as convergence points in the simplex.

Figure 3 (top) shows such a simplex visualization for one participant with the asymmetric progress-velocity momentum circuit model fitted to behavioral choices. The fitted flow fields show policy dynamics biased toward the previously chosen option, thereby inducing persistence. The flow toward the previously chosen goal becomes stronger in high-progress contexts. We repeated the same analyses used to quantify the point-attractor score, this time estimating the score in simplex policy space. Figure 3 (bottom) shows a significant increase in point-attractor scores for flow fields derived from high-progress contexts compared with low-progress contexts, indicating a progress-modulated shift in the dynamic landscape guiding policy transitions. We thus show a circuit-level model of goal selection that can induce persistence through dynamic modes in policy space, with dynamics shaped by environmental rewards, goal progress, progress velocity, choice history, and choice feedback.

**Figure 3:**
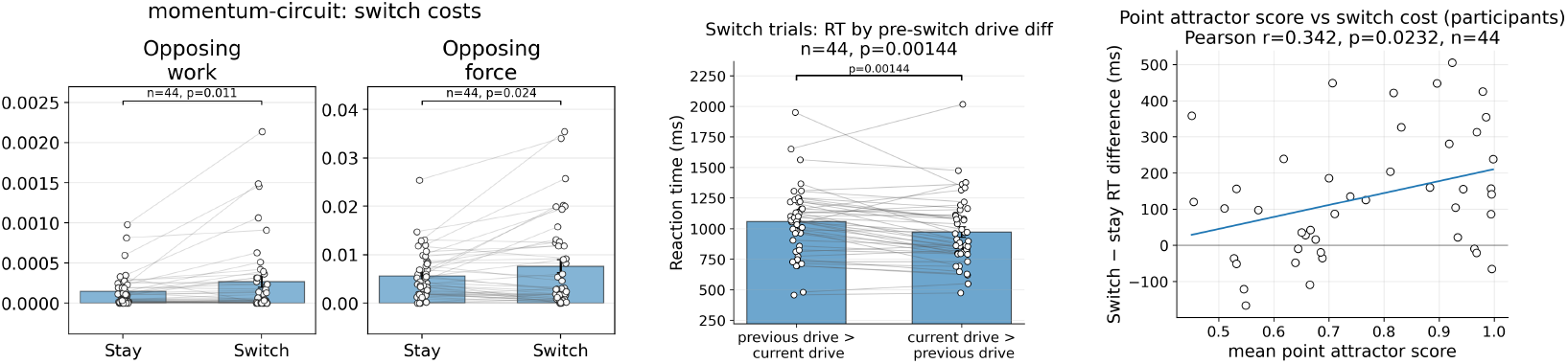
Switching costs as resistance to intrinsic dynamics: (left) Work done opposing the intrinsic dynamics and force opposing the state update are significantly greater in the switch condition than in the stay condition in the latent dynamics of momentum-circuit model, accounting for the emergence of switching costs. (center) Reaction time for switch trials is significantly greater when the previous goal drive is greater than the current goal drive compared to when the current goal drive is greater than the previous goal drive. (right) Participants’ switching cost is positively correlated with their point attractor score, indicating that individual differences in behavioral switching are explained by differences in the dynamical stability of goal policies.

### Ablation experiments

We performed ablation experiments on the momentum-circuit model to understand the contribution of each component to the model’s performance. We ablated the following terms one at a time: progress (*β*_*prog*_), progress velocity (*β*_vel_), interaction (*β*_mom_), context vector (*β*_ctx_), and velocity asymmetry (collapsing 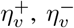). Table 1 shows the results of the ablations evaluated on the validation dataset. We report the model’s negative log-likelihood of choices (val NLL) as well as its performance on key behavioral measures. We evaluated mean-squared errors (MSE) on three behavioral metrics corresponding to the plots shown in Figure 3 (bottom): conditional switching probabilities *P* (switch | *a*_*t* −1_) for each previous action, progress-conditioned switching 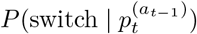 across low, medium, and high progress bins, and dominant-goal choice probability 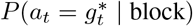 across block conditions (80-20, 70–30, 60–40). The full momentum-circuit model successfully recovers these behavioral metrics, as validated through simulations (details in Appendix Section E2, Appendix Figure 8). The full model performs better in val NLL scores as well as aggregate MSE scores on behavioral measures. Moreover, the momentum-circuit model outperforms the behavior-cloned RNN and the momentum model described in [2] across all metrics.

**Table 1:**
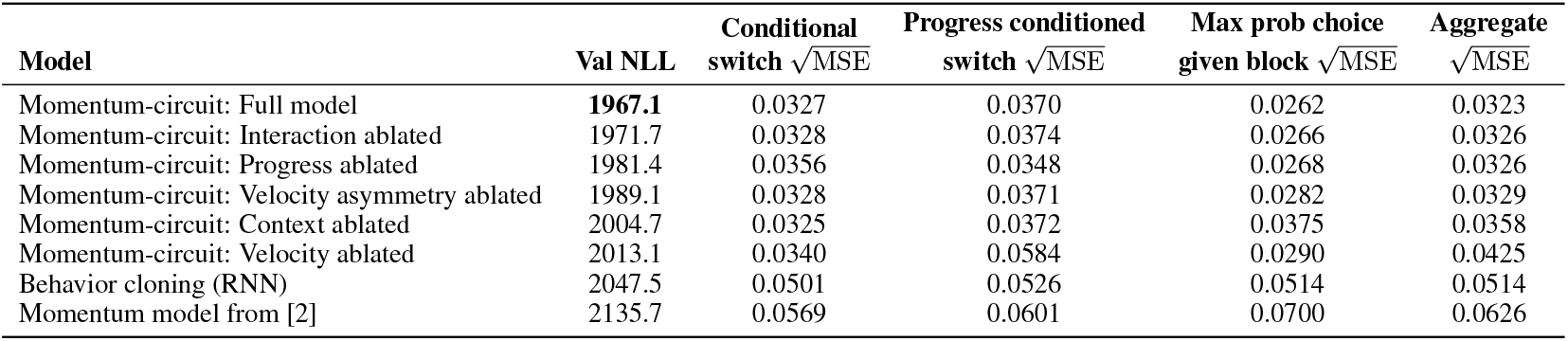
Ablation study of the momentum-circuit architecture on held-out validation data. Performance is evaluated using validation negative log-likelihood (val NLL) and several switch-prediction metrics 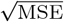.

## 5 Switching Costs as Resistance to Intrinsic Dynamics

We formalize switching costs [25] as resistance to the intrinsic flow field of the learned dynamical system. Let *x*_*t*_ ∈ Δ^2^ ⊂ ℝ^3^ denote the goal state. The model induces a velocity field: *v*(*x*_*t*_) = *π*(*x*_*t*_) −*x*_*t*_, where *π*(*x*_*t*_) is the policy implied by the circuit. Given an empirical trajectory *x*_*t*_ → *x*_*t*+1_, we define the observed displacement: Δ*x*_*t*_ = *x*_*t*+1_ − *x*_*t*_.

We measure how much this movement opposes the intrinsic dynamics via projection onto the velocity field. The instantaneous opposing force is defined as

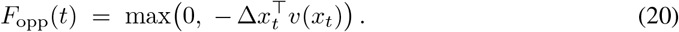

This quantity is zero when the trajectory follows the flow and positive when it moves against it. We define the total opposing work along a trajectory as *W*_opp_ = Σ_*t*_ *F*_opp_(*t*), which captures the cumulative resistance required to deviate from the system’s natural dynamics.

To quantify switching cost, we partition timesteps into stay (*a*_*t*+1_ = *a*_*t*_) and switch (*a*_*t*+1_ ≠ *a*_*t*_) events, and compute the mean opposing work in each condition. Switching cost emerges when: 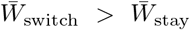 , indicating that switching requires greater work against the intrinsic flow field (Figure 3). When 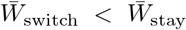 , where the drive to switch is stronger than staying, switching costs do not emerge. This explains why participants often revisit high-progress goals after switching away from them [2], and why some task switches are easier than others [26, 27]. Figure 3 (center) shows that participants’ reaction times in switch trials are significantly higher when the previous goal’s drive exceeds the current goal’s drive, indicating higher switching costs when working against the intrinsic dynamics of the system. Participants’ switching costs are also positively correlated with their mean point-attractor score (Pearson correlation *R* = 0.34, *p* = 0.02), indicating that individual differences in switching are explained by differences in the dynamical stability of goal policies.

## 6 Discussion

Human goal pursuit is a mix of stability and flexibility, stability to ensure goals make consistent progress toward their targets, and flexibility to facilitate switching to better alternatives when available. Previous work shows that human goal commitment is significantly modulated by goal progress, where decision-making vigor increases closer to goal resolution, and by goal progress rates, where repeated failures increase the drive to switch away from the goal [2, 3, 28, 29]. In a normative account, we show that simply incorporating the objective of optimizing discounted returns is sufficient to elicit human-like persistence in RNN actor-critic models whose latent spaces show clear gradations of goal progress and progress rates. We then investigated the latent-space dynamics of human goal commitment by cloning behavioral policies with RNNs. By analyzing velocity flow fields in latent space, we show the emergence of point-attractor-like dynamic modes whose “pointiness” is modulated by goal progress. This suggests a shifting of the energy landscape of neural activity patterns toward sustaining stable goal states in high-progress contexts. We quantified metrics for estimating point-attractor scores that could serve as estimates of the degree of goal commitment.

We introduced a circuit-level mechanistic account of goal commitment alongside a descriptive benchmark of RNN behavior cloning. We incorporated a recurrent competition network motif with self-excitation and cross-inhibition that is modulated by a momentum-generating drive circuit. Momentum signals are computed from the interaction between current progress and a temporally smoothed estimate of recent progress change, and we show that it reasonably approximates time-to-goal-completion, an estimate of goal value for agents optimizing discounted future returns. At the implementational level, we envision a circuit where the goal-progress signal is compared to a time-delayed version to compute progress differences, which are passed through an asymmetric leaky integrator to estimate progress velocity. The resulting velocity signal is then multiplied with the current progress signal to drive progress-dependent escalation of commitment. We show that this recurrent circuit produces similar point-attractor-like dynamic modes in its policy landscape, with higher point-attractor scores in high-progress goal contexts. We show that this model reproduces participants’ behavioral patterns better than the behavior-cloning RNN and simpler behavioral models from previous studies [2]. We further performed ablation experiments on this model to ensure all its components contribute to explaining human behavior. Nevertheless, the mechanistic account put forth in this study is fairly limited, lacking precise neural correlates of momentum computations. A likely candidate is the prefrontal cortex, which is known to use grid-like computations that track goal progress and progress rates in abstract goal navigation, just as they compute spatial progress and velocity during spatial navigation [30].

One of the key interpretations of this dynamical-systems perspective on goal commitment is the origin of switching costs [31, 32]. We claim that switching costs appear not as a consequence of switching goals, but because of the resistance that the intrinsic dynamics of the system offer to state updates. This resistance is significantly higher while switching goals, though not always. At times, switching goals align with the intrinsic dynamics, where staying with the goal is more costly. This accounts for why it is difficult to maintain goal commitment in the presence of distracting tasks, where switching is more amenable to the intrinsic dynamics of the system.

One of the major limitations of this study is that we cannot establish whether the attractor modes are manifestations of behavioral commitment or causally sustain commitment through the mechanistic process we describe. This remains promising future work [33]. Moreover, we studied extended goals in a simplistic setting, whereas naturalistic goal pursuit is far more complex and influenced by several factors other than goal progress and progress rates. Most goals are also hierarchical and compositional, where the complexity involved in memory retrieval and management influences goal selection and maintenance. These limitations form the basis for fresh investigations in this area, which is ripe for new insights into goal-directed agency in the real world, a question pertinent to both natural and artificial intelligence.

## A Task Environment

We consider a three-goal sequential decision-making task. At each time step *t*, the agent selects an action *a*_*t*_ ∈ {0, 1, 2} corresponding to pursuing one of three goals: *cat, hat, car*.

Each goal *i* maintains a progress state 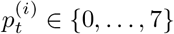 , where 7 is the number of tokens required to complete the goal. Upon completion 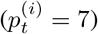, the agent receives a reward *R*_goal_ and the progress resets: 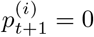 .

At each step, a token is obtained with probability determined by the current block:

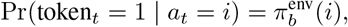

**Figure 4:**
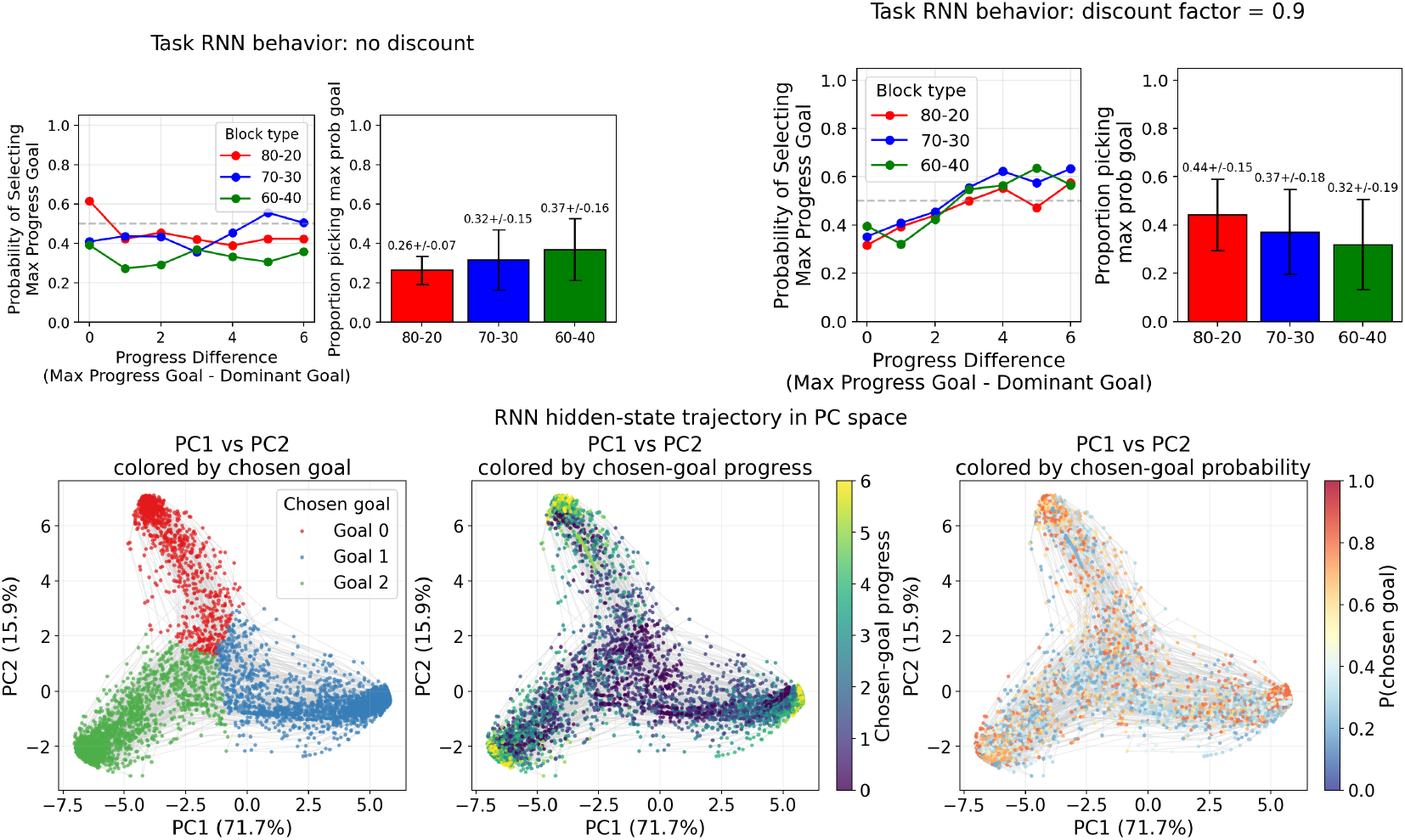
(top)(left) RNN actor-critic models trained on the task show no progress preference when rewards are not discounted. (right) RNN actor-critic models reproduce human block sensitivity and progress preferences when rewards are discounted. (bottom) Latent state of the RNN model (with discounted rewards) visualized in PC space. (panel 1) colored by chosen goal. (panel 2) colored by goal progress. (panel 3) colored by goal progress rate.

where *b* indexes the current block. Blocks last for a fixed number of steps and define reward probability structures (e.g., 80-20, 60-40).

The reward function is:

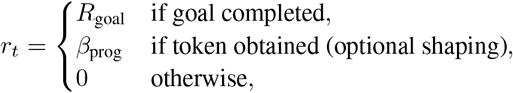

The observation *o*_*t*_ consists of:

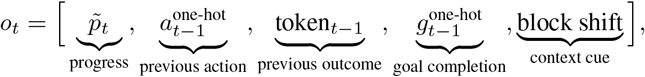

## B Recurrent Policy Architecture

The agent is parameterized by a recurrent actor–critic network. Observations are first embedded:

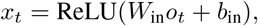

and passed through a gated recurrent unit (GRU):

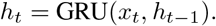

The policy (actor) and value (critic) are defined as:

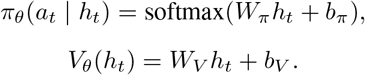

The recurrent hidden state *h*_*t*_ enables integration over past observations and actions, allowing the policy to represent latent goal states and environmental context.

### Trajectory Collection

Training proceeds by sampling trajectories *τ* = (*o*_1_, *a*_1_, *r*_1_, … , *o*_*T*_) from the environment. Actions are sampled from the policy:

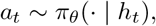

with additional *ϵ*-greedy exploration:

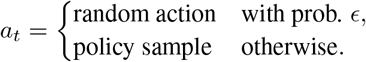

### Return and Advantage Estimation

Discounted returns are computed as: 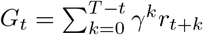

The critic is trained to predict normalized returns: 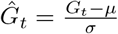,where *µ* and *σ* are running estimates of return mean and variance.

Advantages are computed using the denormalized value estimate: *A*_*t*_ = *G*_*t*_ − *V*_*θ*_(*h*_*t*_)

Advantages are normalized across the trajectory.

### Optimization Objective

The actor loss is:

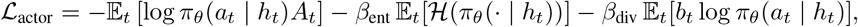

where:

- ℋ is policy entropy (encouraging exploration),
- 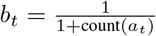is a diversity bonus promoting selection of under-sampled goals.

The critic loss is: 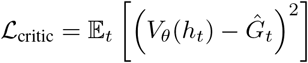

The total loss is: ℒ = ℒ_actor_ + *λ*_*V*_ ℒ_critic_

Parameters are optimized using Adam with gradient clipping.

### Training Procedure

Agents are trained for multiple episodes, each consisting of a sequence of blocks with changing reward structures. At each episode:

1. A trajectory is sampled from the environment.
2. Returns and advantages are computed.
3. Actor and critic losses are evaluated.
4. Parameters are updated via stochastic gradient descent.

Over training, the recurrent dynamics adapt to encode both goal persistence and sensitivity to changing environmental contingencies, giving rise to structured dynamical regimes in hidden-state space.

All analyses were performed on a high-performance computing (HPC) cluster utilizing parallelized workloads distributed across multiple CPU cores.

## C Behavioral Cloning with Recurrent Neural Networks

### Input Representation

For each subject, we construct a single sequence of trials *t* = 1, … , *T*. Each observation *o*_*t*_ ∈ℝ^11^ consists of:

- normalized goal progress *p*_*t*_ ∈ ℝ3,
- previous action encoded as a one-hot vector onehot(*a*_*t*−1_) ∈ {0, 1}^3^,
- previous reward outcome *r*_*t*−1_ ∈ {0, 1},
- previous goal completion indicator *g*_*t*−1_ ∈ {0, 1}^3^,
- block boundary indicator *b*_*t*_ ∈ {0, 1}.

**Figure 5:**
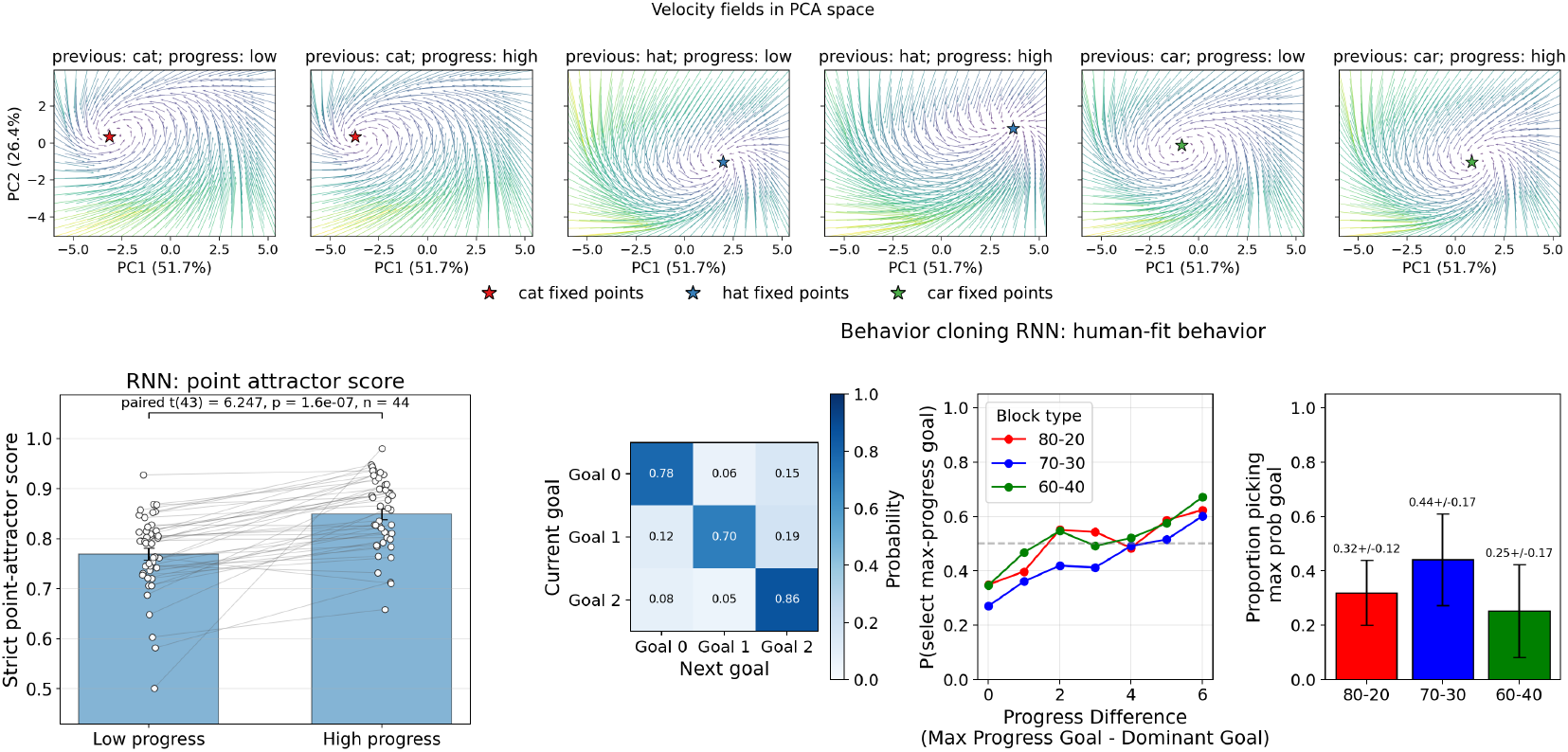
(top) Context-dependent intrinsic velocity fields of behavior-cloned RNN model of a single participant in PC space showing point-attractor-like structure. Each panel shows the velocity field for a specific previous goal and progress level. (bottom) (left) Significantly higher point-attactor scores in high-progress contexts compared to low-progress contexts showing attractor-structure evolution with environmental context. (right) Behavioral cloning through RNN models reproducing human persistence patterns and progress preferences.

Thus,

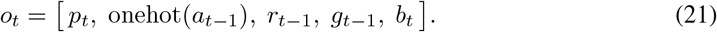

Actions are discrete choices *a*_*t*_ ∈ {0, 1, 2}.

### Model Architecture

We use a recurrent neural network with hidden state *h*_*t*_ ∈ ℝ^*H*^:

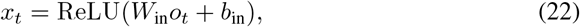

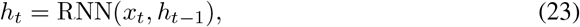

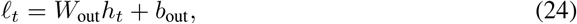

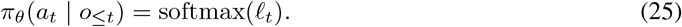

### Training Objective

We fit parameters *θ* via maximum likelihood estimation:

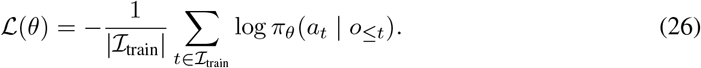

Optimization is performed using Adam with gradient clipping. Early stopping is applied based on validation negative log-likelihood.

### Data Splitting

Trials are randomly partitioned into disjoint sets:

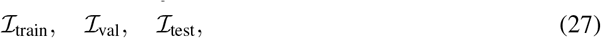

with proportions 75%, 12.5%, and 12.5%, respectively. Splitting is interspersed across time to preserve coverage of the full behavioral sequence.

### Evaluation Metrics

We evaluate performance on held-out trialsℐusing:

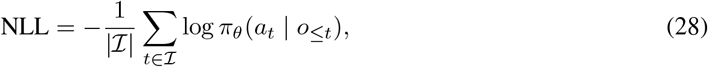

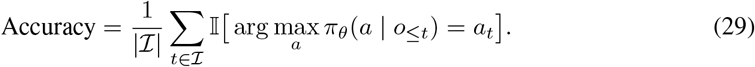

To assess agreement in marginal choice structure, we compute:

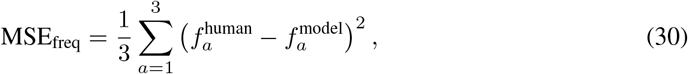

where *f*_*a*_ denotes empirical choice frequencies.

### Hidden State Extraction

After training, we compute the hidden trajectory 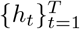 under the fitted model. These latent states are used for subsequent dynamical systems analyses (e.g., vector fields and attractor structure).

### Per-Subject Fitting

All models are fit independently per subject, yielding parameters *θ*^(*s*)^. Group-level statistics are obtained by aggregating performance metrics across subjects.

### C.1 Intrinsic Velocity Field Analysis

### Hidden State Extraction and Dimensionality Reduction

For each subject, we compute the hidden state trajectory 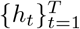 from the fitted RNN. We perform principal component analysis (PCA) on {*h*_*t*_} and obtain a low-dimensional embedding:

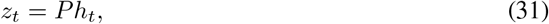

where *P* ∈ ℝ^*d*×*H*^ contains the top 2 principal components.

### Context Definition

We define discrete behavioral contexts based on:

- previous goal identity *g* ∈ {0, 1, 2},
- goal progress level (low vs. high).

Formally, a context *c* = (*g*, bin) selects timepoints satisfying:

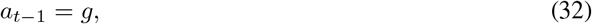

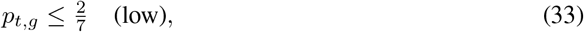

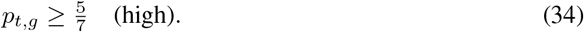

For each context, we compute a prototype observation:

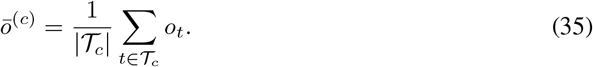

### Intrinsic Velocity Field

We define the intrinsic dynamics by evaluating the RNN transition from arbitrary latent states under a fixed context:

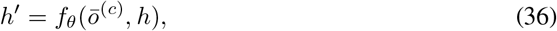

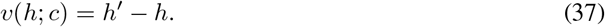

We construct a grid {*z*_*i*_} in PCA space and map each point back to latent space:

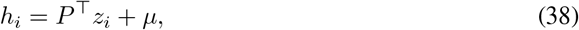

where *µ* is the PCA mean.

The velocity field is then computed as:

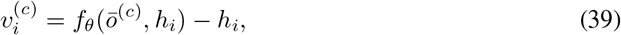

and projected back to PCA coordinates:

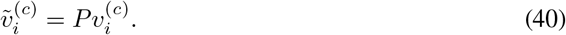

The projected field 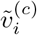 is visualized as a quiver plot over the PCA grid: 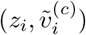. Velocity magnitude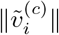 is used for color coding.

### Fixed Point Estimation

We estimate candidate fixed points as local minima of the velocity magnitude:

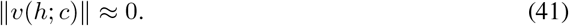

Specifically:

1. Compute speeds 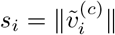.
2. Select grid points below a quantile threshold.
3. Identify local minima in the grid neighborhood.
4. Cluster nearby minima using a distance threshold. Cluster centers define candidate fixed points:

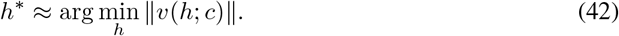

## D Strict Point-Attractor Metric and Synthetic Validation

### D.1 Definition of the Strict Point-Attractor Metric

To quantify the presence of a true point attractor in a vector field, we define a strict metric that combines multiple necessary dynamical properties. The goal is to assign a high score only when a candidate point exhibits inward flow, basin support, and full local stability.

#### Candidate selection

Candidate points are identified as locations where the vector field magnitude is low. Specifically, we select grid points whose speed lies below a chosen quantile threshold, with an additional constraint that selected candidates are spatially separated.

#### Inward convergence

For each candidate point *x*_0_, we measure whether nearby vectors point toward it. For neighboring points *x*, let *v*(*x*) denote the vector field. The inward convergence score is defined as:

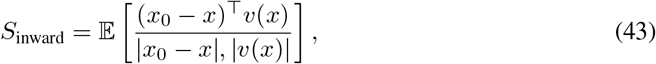

where the expectation is taken over a local neighborhood of *x*_0_.

This quantity reflects whether the local flow is directed toward the candidate point.

#### Basin support

To assess whether the candidate has a meaningful basin of attraction, we evaluate short trajectories initialized in a surrounding region. For each initial point *x*, we compute the change in distance to the candidate after a fixed number of steps:

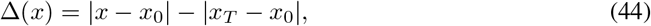

where *x*_*T*_ is obtained by iterating the dynamics.

We define the basin score as a combination of:

- the fraction of points with Δ(*x*) *>* 0, and
- the fraction of points that enter a local neighborhood of *x*_0_.

This captures whether trajectories converge toward the candidate beyond its immediate vicinity.

#### Point-specific stability

We assess local stability using the Jacobian of the vector field:

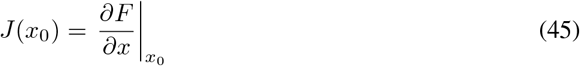

Let *λ*_*i*_ denote the real parts of the eigenvalues of *J*(*x*_0_). We define a point-specific stability score as:

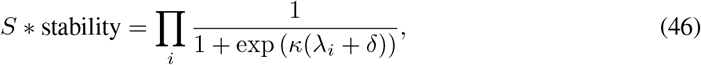

where *κ >* 0 controls sharpness and *δ >* 0 enforces a margin.

This formulation penalizes both unstable directions (*λ*_*i*_ *>* 0) and neutral directions (*λ*_*i*_ ≈0), ensuring that line attractors do not receive high scores.

#### Final metric

The strict point-attractor score is defined as:

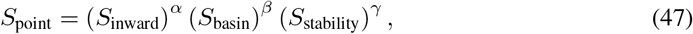

where *α* = 0.35, *β* = 0.16, *γ* = 0.48 are weighting exponents chosen to maximize the score’s ability to distinguish point attractors from other dynamical regimes.

This multiplicative form ensures that all components must be simultaneously present for a high score.

### D.2 Synthetic Validation

To validate the metric, we constructed a set of synthetic two-dimensional dynamical systems with known ground-truth structure. Each system defines a vector field:

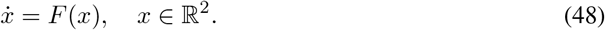

We generated multiple classes of systems:

- **Point attractors:** Stable fixed points were generated using perturbed linear contraction dynamics:

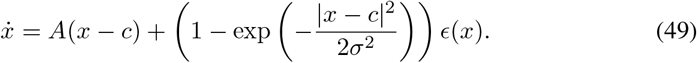

Here, *c* is the attractor location, *A* has negative eigenvalues, and *ϵ*(*x*) is a smooth perturbation that vanishes near *x* = *c*.
- **Limit cycles:** Limit cycles were generated using radial stabilization and tangential rotation:

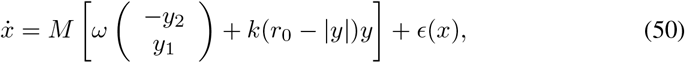

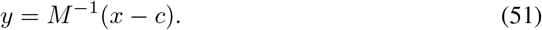

Here, *r*_0_ is the target radius, *k* controls radial attraction, and *ω* sets angular velocity.
- **Line attractors:** Line attractors were generated by separating dynamics along tangent and normal directions:

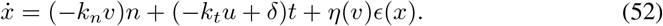

The scalar coordinates are:

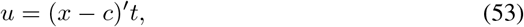

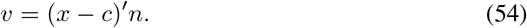

Here, *k*_*n*_ *>* 0 enforces contraction toward the line, while *k*_*t*_ is close to zero, giving approximately neutral dynamics along the line.
- **Repellers:** Repellers were generated as unstable linear systems:

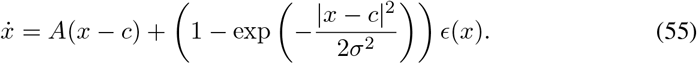

Here, *A* has positive eigenvalues, producing outward flow in all directions.
- **Saddles:** Saddles were generated using one unstable and one stable direction:

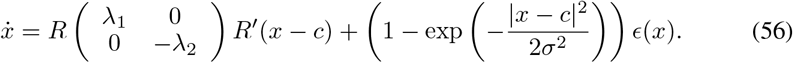

Here, *λ*_1_ *>* 0 and *λ*_2_ *>* 0.
- **Random flows:** Random flows were generated as smooth Fourier vector fields:

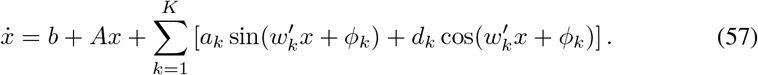

The coefficients, frequencies, and phases were sampled randomly, producing smooth dynamics without a planted attractor.

Across all systems, parameters such as location, strength, anisotropy, and rotation were randomized. In addition, smooth spatial perturbations were added to introduce variability while preserving the qualitative structure of each dynamical class.

**Figure 6:**
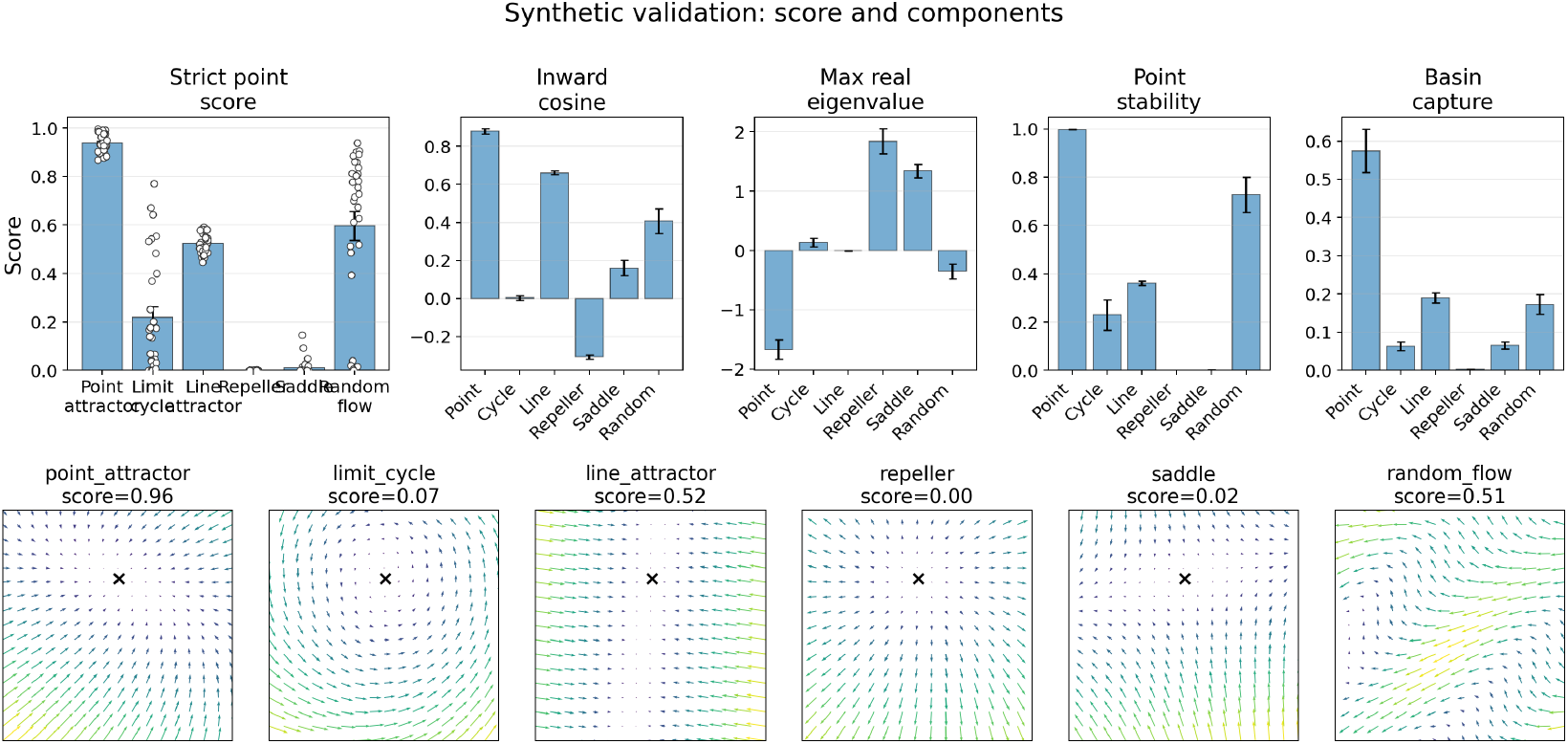
Synthetic validation of the strict point-attractor score: (top) Summary of point attractor scores by dynamical system class. (bottom) Example velocity fields and their corresponding point attractor scores.

#### Evaluation procedure

For each synthetic field:

1. Candidate points were identified using low-speed regions.
2. Each candidate was scored using the strict metric.
3. The highest-scoring candidate was selected.

This procedure was repeated across multiple randomized replicates for each class.

#### Validation criterion

A valid metric should assign:

- high scores to point attractors,
- low scores to limit cycles and random flows,
- reduced scores to line attractors due to neutral directions,
- low scores to repellers and saddles due to instability.

The strict metric reliably distinguished point attractors from all other dynamical regimes. In particular, the point-specific stability component was critical in separating point attractors from line attractors, which otherwise exhibit local convergence but lack a unique stable equilibrium.

These results demonstrate that the metric captures the defining properties of point attractors and is not driven by artifacts such as slow dynamics or tangential flow (see appendix Figure 6).

## E Momentum-Circuit Model

### State and dynamics

The model maintains a latent goal state:

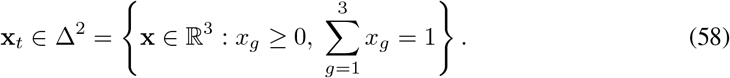

### Recurrent circuit

The recurrent interaction matrix *W* ∈ ℝ^3×3^ is defined as:

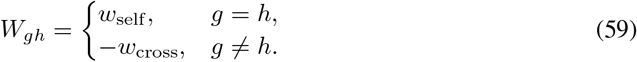

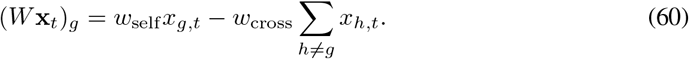

**Figure 7:**
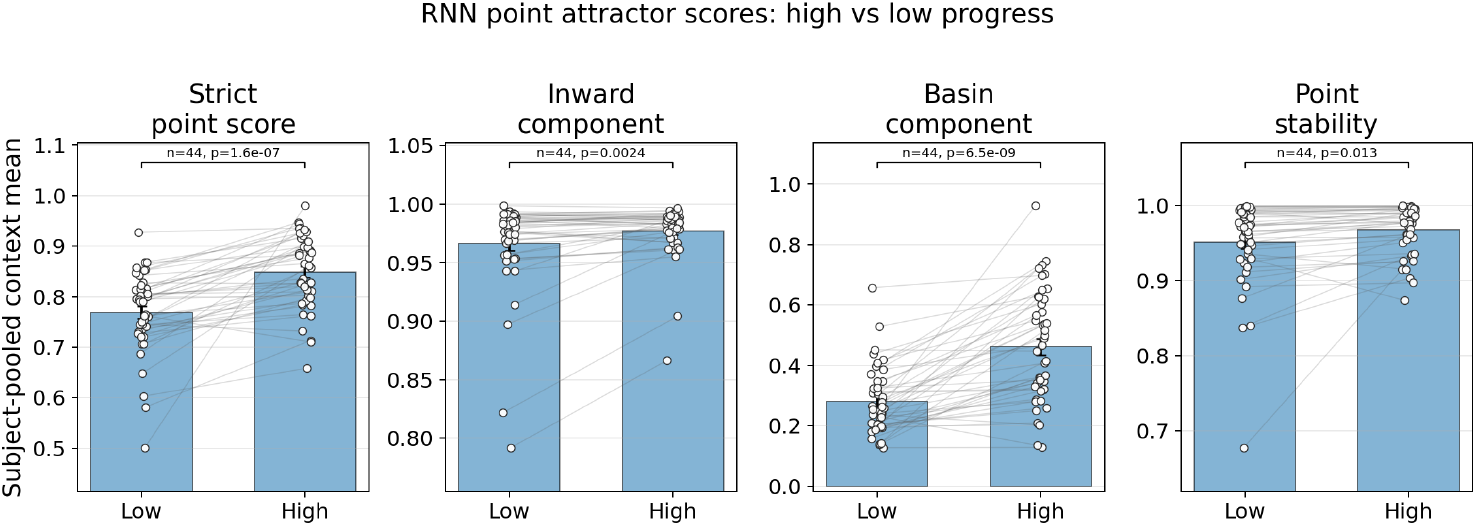
Significantly higher point-attactor scores in high-progress contexts compared to low-progress contexts showing attractor-structure evolution with environmental context in Behavior-cloned RNN model.

### Asymmetric momentum computations

Let **p**_*t*_ ∈ [0, 1]^3^ denote normalized progress. First define instantaneous progress change:

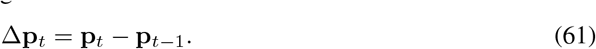

We separate positive and negative progress changes:

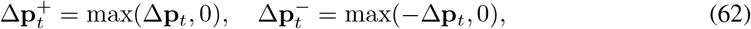

where the max is taken elementwise.

Velocity is computed via a leaky integration with asymmetric gains:

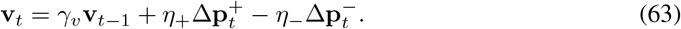

This allows positive progress and negative (or stalled) progress to have distinct impacts on the velocity signal.

### Drive and policy

The total drive now includes additional inputs:

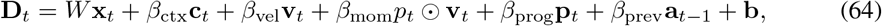

where:

- **p**_*t*_: direct progress input
- **a**_*t*−1_: one-hot vector of the previous action

Choice probabilities follow:

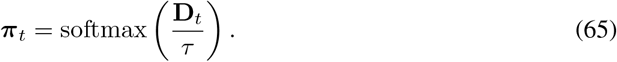

### State update with choice feedback

The latent state evolves as:

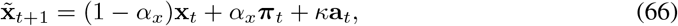

where a_*t*_ is the chosen action (one-hot) and *κ* is the choice feedback gain.

The state is then renormalized onto the simplex:

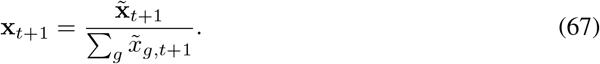

The model decomposes goal commitment into:

- **Recurrent dynamics:** *W* **x**_*t*_ defines attractor structure.
- **Asymmetric velocity: v**_*t*_ distinguishes gains from losses in progress.
- **Momentum: p**_*t*_ ⊙ **v**_*t*_ amplifies sustained success.
- **Direct progress drive: p**_*t*_ biases toward advanced goals.
- **Action persistence: a**_*t*−1_ and **a**_*t*_ reinforce recent choices.

This yields a dynamical system where persistence arises from both attractor structure and history-dependent progress signals, while switching reflects competition between accumulated evidence streams.

### E.1 Momentum as an Approximation of Time-to-Goal Completion

Let *x*(*t*) ∈ [0, 1] denote fractional progress toward a goal, with *x* = 1 corresponding to goal completion, and let 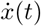 denote the rate of progress. We define a momentum-like quantity:

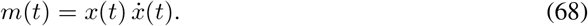

The exact remaining time to goal completion is given by:

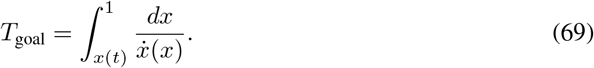

Under a local constant-velocity approximation, where 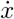 is assumed to vary slowly over the remaining trajectory, this simplifies to:

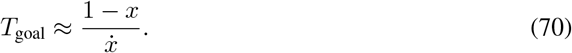

Substituting 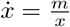yields:

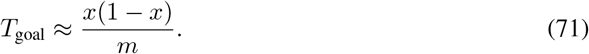

Thus, momentum is inversely related to time-to-goal completion, up to a state-dependent scaling factor:

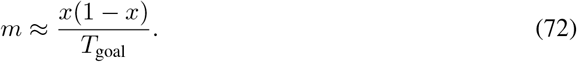

This relationship shows that *m* is not a direct estimator of time-to-goal, but rather a *progress-weighted inverse time-to-completion*. The multiplicative factor *x*(1− *x*) introduces a characteristic dependence on the current state: it is maximal at intermediate progress (*x* ≈ 0.5) and vanishes near the start and end of the trajectory. This could potentially explain human motivation patterns pertaining to goal pursuit, where they lose motivation at the last stretch of goal-directed activity, wherein the drive to finish the last 1% is substantially lower than the drive mid-progress.

Consequently, *m* provides a locally accurate proxy for inverse time-to-goal in mid-progress regimes, while systematically underestimating it near boundary states. Despite this limitation, the quantity 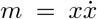 captures a behaviorally meaningful signal, reflecting the interaction between current commitment (progress) and ongoing motion (rate of progress), and thus serves as a compact summary of goal-directed dynamics.

### E.2 Behavior Recovery Analysis

To complement parameter recovery, we evaluated whether the model can reproduce *behavioral statistics* of synthetic agents generated from known ground-truth parameters. This provides a functionally relevant validation, as different parameter configurations can produce similar observable behavior.

#### Synthetic Data Generation

For each recovery simulation, we sampled ground-truth parameters from predefined ranges and instantiated the momentum-circuit model. Using the empirical task structure (progress and context sequences from a human subject), we generated synthetic choice sequences by rolling out the model policy:

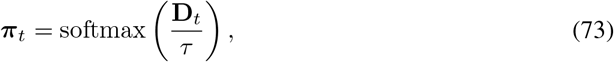

**Figure 8:**
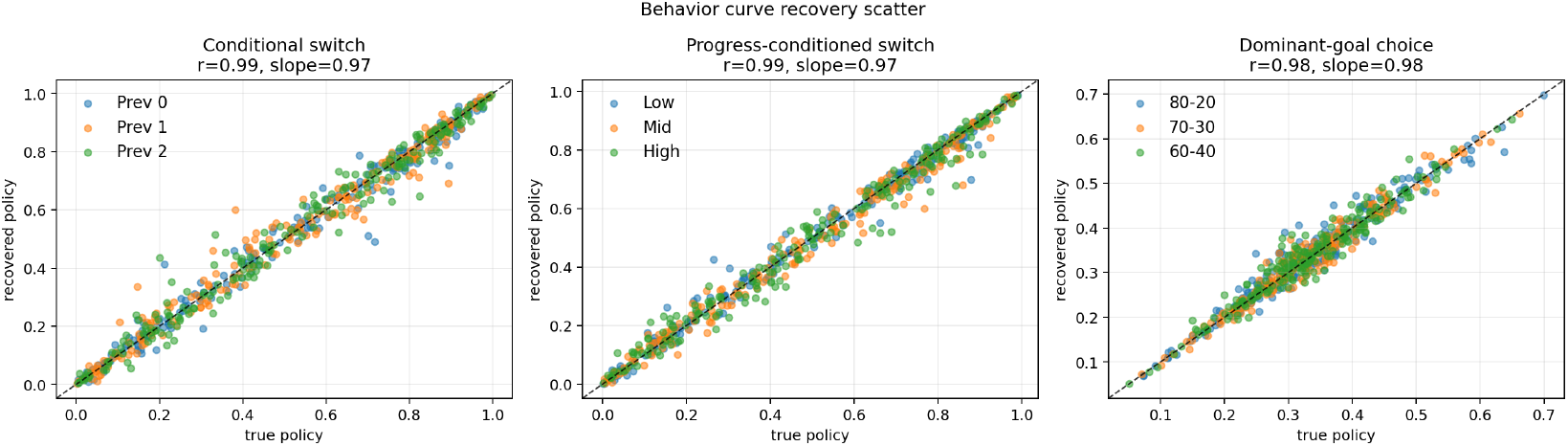
Behavioral recovery of structured decision statistics across simulations. Each panel compares ground-truth (x-axis) and recovered (y-axis) values computed from the fitted model’s stochastic policy. Left: conditional switching probabilities *P* (switch | *a*_*t−*1_) for each previous action. Middle: progress-conditioned switching 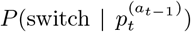 across low, medium, and high progress bins. Right: dominant-goal choice probability 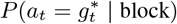 across block conditions (80-20, 70–30, 60–40). Each point corresponds to one simulation; colors denote conditioning levels. Dashed lines indicate identity.

where the drive is given by:

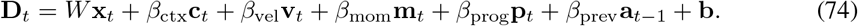

Actions were sampled stochastically from this policy to preserve realistic variability. To ensure identifiability, we rejected degenerate simulations with low entropy or highly imbalanced action frequencies.

#### Behavioral Metrics

We summarized behavior using task-relevant conditional statistics:

#### Dominant-goal choice probability (by block)

For each block condition *b* ∈ {80-20, 70-30, 60-40 }, we compute the probability of selecting the dominant goal:

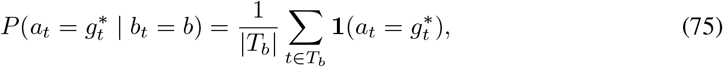

where 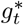 denotes the dominant (highest-reward-probability) goal on trial *t*, and *T*_*b*_ is the set of trials belonging to block *b*.

#### Conditional switching by previous action

We quantify action-dependent persistence by computing:

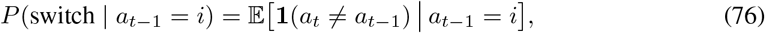

for each action *i* ∈ {1, 2, 3}.

#### Progress-conditioned switching

We measure how switching depends on goal completion progress:

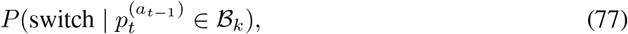

where 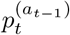 denotes the progress of the previously selected goal, and ℬ_*k*_ are discrete progress bins (low, medium, high).

#### Recovery Procedure

For each synthetic dataset, we refit the model using the same optimization procedure as in the main analysis. After fitting, we computed the recovered behavioral statistics using:

- **Stochastic policy estimate:** behavioral metrics computed directly from the model’s predicted action probabilities ***π***_*t*_.
- **Sampled rollout:** behavioral metrics computed from new sequences generated by sampling from the recovered policy.

### Behavior Recovery Evaluation

For each synthetic dataset, we fit the model to the generated choice sequence using the same optimization procedure as in the main analysis. We then evaluated whether the recovered model reproduced the *behavioral structure* of the synthetic agent by comparing behavioral statistics computed from the recovered stochastic policy to those computed from the ground-truth generating policy. Specifically, recovered behavioral metrics were computed directly from the model’s predicted action probabilities ***π***_*t*_, conditioned on the same task covariates and synthetic action history used during generation, rather than from deterministic rollouts.

Behavior recovery focused on three task-relevant statistics: (1) conditional switching probabilities given the previous action, (2) progress-conditioned switching probabilities across low, medium, and high progress states, and (3) dominant-goal choice probabilities across block conditions. Recovery quality was quantified across simulations using Pearson correlation, linear regression slope, and root mean squared error (RMSE) between ground-truth and recovered behavioral statistics. Successful recovery therefore indicates that the fitted model reproduces the functional mapping from task state variables (progress, context, and action history) to action-selection behavior, rather than merely matching individual parameter values.

## F Simplex Landscape Analysis

### Simplex Representation

The goal state lies on the simplex:

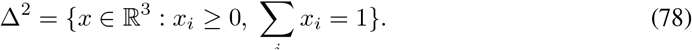

We embed *x* into 2D coordinates for visualization using barycentric projection.

### Context Anchoring

For each condition *c* = (previous goal, progress bin), we construct anchor variables by averaging over trials matching the condition:

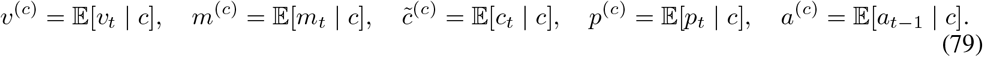

Here, *v*_*t*_ and *m*_*t*_ denote velocity and momentum, *c*_*t*_ is the inferred context, *p*_*t*_ is the progress vector across goals, and *a*_*t*−1_ is the previous action (one-hot encoded).

These anchors define a fixed input condition under which the dynamical system is evaluated.

### Vector Field Definition

The drive is given by:

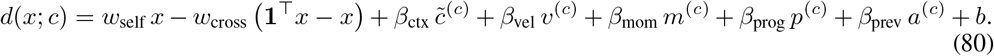

The policy is:

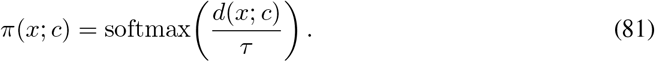

The induced vector field is:

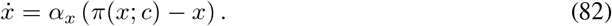

To ensure the dynamics remain within the simplex, we project onto the tangent space:

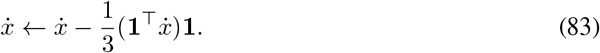

#### Numerical Integration

We simulate trajectories using Euler updates:

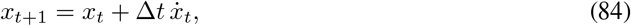

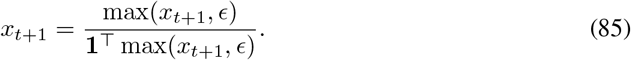

#### Grid Construction

We discretize the simplex using a triangular grid:

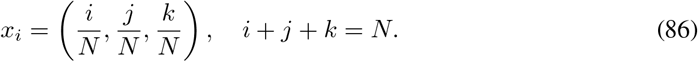

**Figure 9:**
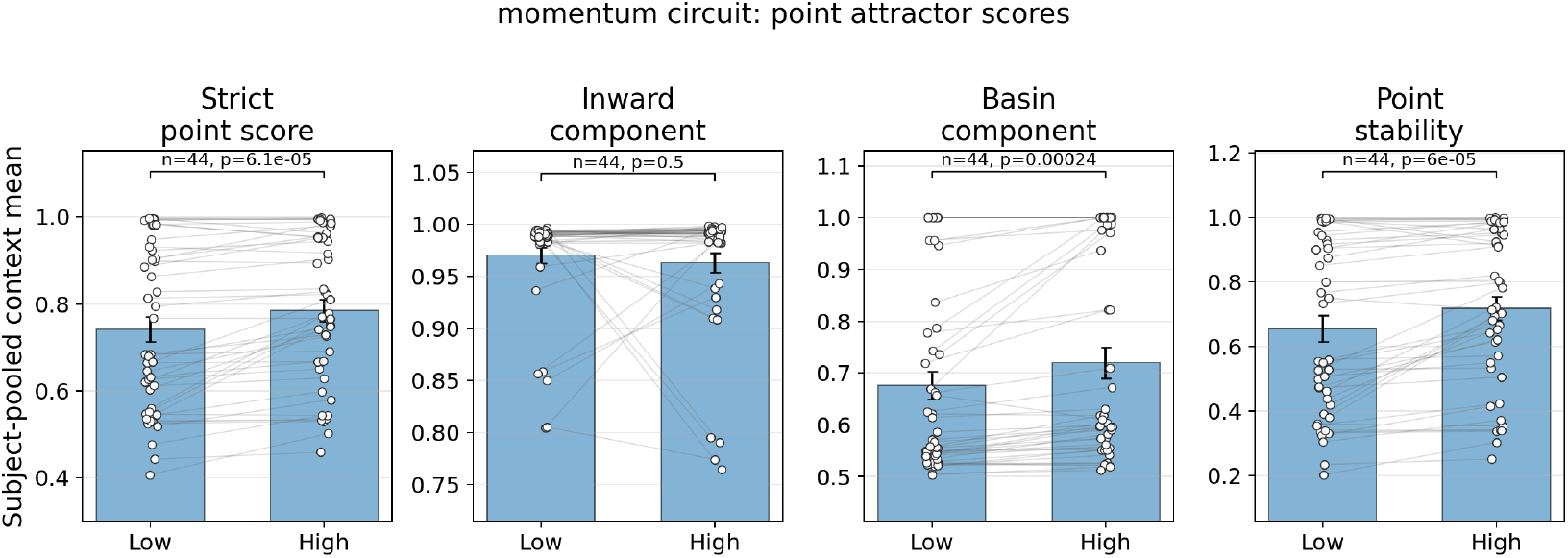
Significantly higher point-attactor scores in high-progress contexts compared to low-progress contexts showing attractor-structure evolution with environmental context for the momentum-circuit model.

The vector field is evaluated at each grid point. We visualize the projected vector field 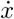(quiver plot), the scalar field log 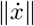, and trajectories from random initial conditions.

#### Attractor Detection

We simulate trajectories 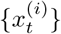 from random initial conditions and extract endpoints:

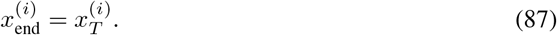

Endpoints are clustered using a distance threshold *ϵ*:

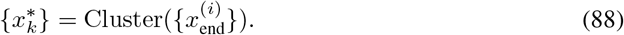

The resulting landscape characterizes the number and location of attractors, basin geometry, and convergence strength via 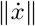. Differences across conditions reveal how velocity, momentum, progress, contextual signals, and action history jointly shape the stability and geometry of goal representations.

## G Strict Point-Attractor Analysis for the Momentum-Circuit Model

We apply the same strict point-attractor metric described in the appendix section D to the momentum-circuit model. Here we describe how the underlying dynamical system differs and how the metric is instantiated in this setting.

### Dynamical System on the Simplex

Unlike the RNN, whose dynamics evolve in a high-dimensional latent space, the momentum-circuit model defines an explicit dynamical system over the goal simplex:

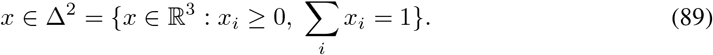

For a fixed behavioral context *c*, the vector field is given analytically by:

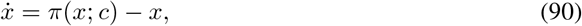

where *π*(*x*; *c*) is the softmax policy induced by the model drive.

Thus, the momentum-circuit model provides a closed-form vector field, rather than requiring rollout-based estimation as in the RNN.

### Context Conditioning

Contexts are defined identically to the RNN analysis, using:

- previous goal identity,
- goal progress (low vs. high).

For each context *c*, we compute anchor variables by averaging over trials matching the condition:

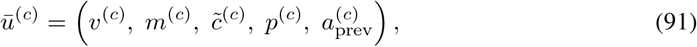

where

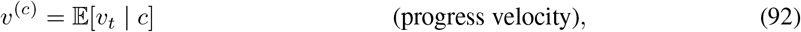

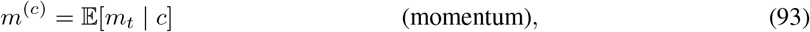

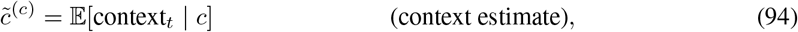

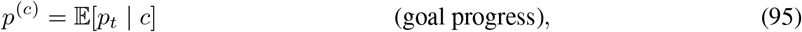

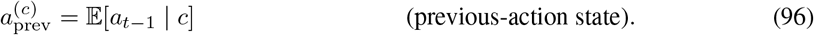

These anchors define a fixed input condition under which the vector field is evaluated:

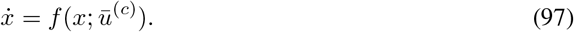

### Candidate Fixed Points

As in the RNN analysis, candidate fixed points are identified as low-speed regions of the vector field evaluated on a simplex grid:

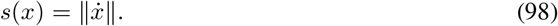

Points below a quantile threshold are selected and filtered to enforce spatial separation.

### Metric Evaluation

All strict point-attractor components (slowness, inward convergence, radial contraction, local stability, and basin support) are computed exactly as in the RNN case, but now directly on the simplex vector field.

A key difference is that:

- distances and directions are computed in simplex coordinates,
- the Jacobian is evaluated in the simplex tangent plane,
- rollouts are performed using explicit Euler updates with projection onto the simplex.

### Jacobian in the Tangent Plane

To respect the simplex constraint 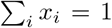, we compute the Jacobian in the tangent subspace:

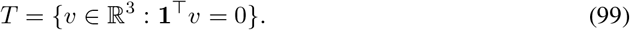

The Jacobian *J* is estimated numerically via finite differences along a basis of *T* , and stability is determined from the maximum real eigenvalue.

### Basin Analysis

Basin support is evaluated by short rollouts:

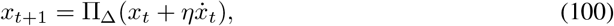

where Π_Δ_ projects onto the simplex. This yields estimates of:

- radial contraction over multiple steps,
- basin capture probability.

### Aggregation and Comparison

Metrics are computed across all contexts and aggregated within subjects:

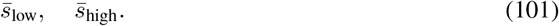

We compare low and high progress using paired tests across subjects, identical to the RNN analysis.

The momentum-circuit model thus provides a mechanistic instantiation of attractor dynamics, allowing direct interpretation of:

- how recurrent competition shapes attractor geometry,
- how momentum (*β*_mom_) stabilizes goal states,
- how progress (*β*_prog_) modulates attractor strength.

Because the vector field is analytically defined, this analysis provides a direct link between model parameters and the emergence of point-attractor structure.

## H Emergence of switching costs

We quantify switching cost as movement against the intrinsic velocity field of the learned dynamical system.

### Velocity field

Let *x*_*t*_ ∈ Δ^2^ ⊂ ℝ3 denote the goal state on the simplex. The model defines a velocity field:

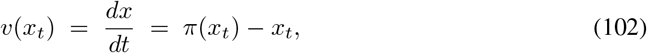

where *π*(*x*_*t*_) ∈ Δ^2^ is the softmax policy induced by the circuit.

### Observed state displacement

Given the empirical trajectory *x*_*t*_ → *x*_*t*+1_, we define:

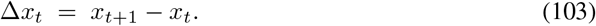

### Alignment with the flow field

We measure alignment between observed movement and intrinsic dynamics:

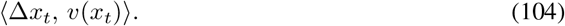

### Opposing force

We define the instantaneous opposing force as:

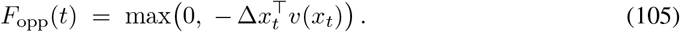

### Accumulated opposing work

Total opposing work along a trajectory is:

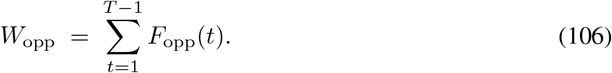

This corresponds to the continuous form:

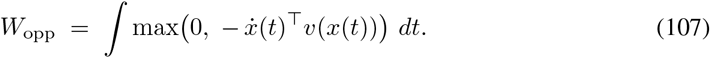

### Stay vs. switch decomposition

We partition timesteps into:

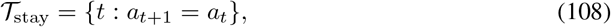

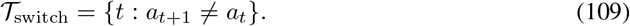

Mean opposing work is then:

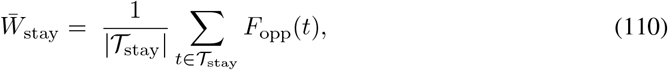

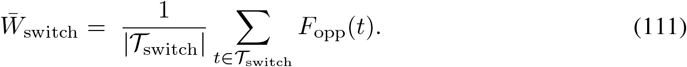

Switching cost emerges if:

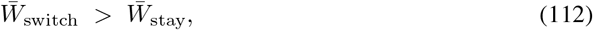

indicating that switching requires greater work against the intrinsic flow field of the dynamical system.

